# Two redundant paralogs of MAR1-BINDING FILAMENT PROTEIN affect B-type granule initiation in wheat

**DOI:** 10.64898/2026.04.21.719982

**Authors:** Nitin Uttam Kamble, Andrés Ortiz, Rokas Kubilinskas, Brendan Fahy, Kay Trafford, David Seung

**Affiliations:** John Innes Centre, Norwich Research Park, Norwich, NR4 7UH, UK; Indian Institute of Science Education and Research, Thiruvananthapuram, India

## Abstract

Starch synthesis in wheat endosperm involves the initiation of large A-type starch granules during early grain development, followed by small B-type granules in later grain development. It is established that MAR-BINDING FILAMENT-LIKE PROTEIN 1 (MFP1) plays an important role in granule initiation in Arabidopsis chloroplasts, but how it influences A- and B-type initiations in wheat amyloplasts is not known. We discovered that due to a gene duplication in cereals, wheat contains two MFP1 paralogs, MFP1.1 and MFP1.2, which are both expressed in the developing endosperm. We generated a series of durum wheat mutants defective in all homoeologs of either MFP1.1 or MFP1.2, or both. While starch granule size distributions and granule morphology of *mfp1.1* and *mfp1.2* mutants were identical to those of the wild-type, the *mfp1.1 mfp1.2* mutants had fewer, but larger B-type granules – suggesting that the two paralogs play redundant roles in B-type granule initiation. Consistent with this, both paralogs interacted with B-GRANULE CONTENT 1 (BGC1), a key protein required for proper B-type granule initiation in wheat, and both paralogs could partially complement defects in starch initiation in the Arabidopsis *mfp1* mutant. Our work demonstrates that MFP1 is required for establishing correct starch granule number in non-photosynthetic amyloplasts, but its role in wheat is limited to B-type granule initiation.

**One-sentence summary:** Wheat has two MFP1 paralogs that interact with the granule initiation protein, BGC1 and influence B-type granule initiation in non-photosynthetic amyloplasts of endosperm.

## Introduction

There is much diversity in starch granule morphology among grass species (Matsushima et al., 2013; Tetlow and Emes, 2017; Watson-Lazowski et al., 2026). Grains of the Triticeae (e.g., wheat, barley and rye) have two distinct types of starch granules, large, lenticular A-type granules and small, spherical B-type granules. These arise from two separate waves of granule initiation, where A-type granules are initiated as early as ∼4 days post anthesis (dpa), while B-type granules are initiated later, at ∼15-20 dpa (Parker, 1985; Bechtel et al., 1990; Chen et al., 2024). A single A-type granule is initiated per amyloplast, whereas multiple B-type granules are thought to initiate at least partly in amyloplast stromules (Parker, 1985; Langeveld et al., 2000; Esch et al., 2023). Understanding how this granule initiation pattern is established is important for improving wheat quality, since starch granule size distributions affect starch physicochemical properties and end-use quality (Lindeboom et al., 2004).

Many of the proteins involved in starch granule initiation were originally discovered and characterised in Arabidopsis leaves (Seung and Smith, 2019; Abt and Zeeman, 2020; Uauy et al., 2025). Arabidopsis chloroplasts contain ∼5-7 starch granules, which are formed within stromal pockets between thylakoid membranes (Crumpton-Taylor et al., 2012; Burgy et al., 2021; Esch et al., 2022). These granules are initiated by the glucosyltransferase, Starch Synthase 4 (SS4) (Roldán et al., 2007), which interacts with PROTEIN TARGETING TO STARCH 2 (PTST2) (Seung et al., 2017). It is proposed that PTST2 binds to maltooligosaccharide primers using its Carbohydrate Binding Module 48 (CBM48) domain, and delivers them to SS4 for further elongation. PTST2 is partly bound to the thylakoid membrane via its interaction with the MAR1-BINDING FILAMENT-LIKE PROTEIN (MFP1) (Seung et al., 2018). MFP1 associates strongly with the stromal side of the thylakoid membranes only (Jeong et al., 2003). MFP1 and PTST2 co-localise in a discrete, punctate pattern, which could represent sites of granule initiation (Seung et al., 2018). Indeed, manipulating MFP1 localisation is sufficient to direct granule initiations to specific locations (Sharma et al., 2024). For example, directing of MFP1 to the inner envelope membrane (via fusions to the N-terminal domain of Tic40) is sufficient to generate starch granules in that aberrant location.

From these discoveries in Arabidopsis, we are now beginning to understand how variation in the functions of these proteins contributes to the diverse initiation patterns observed in non-photosynthetic organs, including the A- and B-type granules of wheat (Uauy et al., 2025). Granule initiation in wheat endosperm requires many of the orthologs of the Arabidopsis initiation proteins, including the PTST2 ortholog, referred to as B GRANULE CONTENT1 (BGC1) (Chia et al., 2020). Both SS4 and BGC1 are important for the control of A-type granule initiation, as knockout of either gene leads to supernumerary initiations during early grain development (Hawkins et al., 2021). However, B-type granule initiation requires the activity of the plastidial Phosphorylase (PHS1), which also interacts with BGC1 (Kamble et al., 2023). Knockout of PHS1 dramatically reduces the frequency of B-type granule initiations, but does not affect A-type granule number, demonstrating that A- and B-type granules initiate via distinct biochemical mechanisms. Interestingly, B-type granule initiation is sensitive to reductions in BGC1 activity (Chia et al., 2020). This was shown through hexaploid bread wheat lines where two BGC1 homoeologs have been knocked out (leaving only one active BGC1 homoeolog), or tetraploid durum wheat lines combining a knockout mutation in one homoeolog and a hypomorphic missense mutation in the other. In both cases, a severe reduction in B-type granule number was observed, with no detectable effects on A-type granules.

Although this important role of BGC1 has been established in wheat endosperm, it is not yet known whether MFP1 orthologs also play an important role in this organ, or in non-photosynthetic organs of plants generally. This is particularly pertinent given that amyloplasts of non-photosynthetic organs do not contain thylakoid membranes like chloroplasts, and thus granules cannot initiate within defined stromal pockets between thylakoids as in chloroplasts. Furthermore, MFP1 was duplicated during grass evolution, leading to two paralogs in cereals (Seung et al., 2018). It is unknown whether these paralogs play similar or distinct roles.

Here, we examined the role of these MFP1 paralogs in granule initiation in wheat endosperm. We found that the two paralogs are fully redundant with respect to each other, both interacting with BGC1, and together playing an important role in B-type granule initiation. This demonstrates the importance of MFP1 in granule initiation in non-photosynthetic amyloplasts.

## Results

### Both paralogs of MFP1 are expressed in the endosperm

To examine the role of MFP1 paralogs in wheat, we first identified the wheat orthologs of Arabidopsis MFP1. Consistent with a duplication of *MFP1* genes in the grasses (Seung et al., 2018), BLASTp searches against the genomes of bread wheat (*Triticum aestivum* L.; IWGSC; Appels et al. (2018)) and durum wheat (*Triticum turgidum* L.; Svevo 1.1; Maccaferri et al. (2019)) revealed two sets of homoeologs. One set encoded on group 1 chromosomes we refer to as MFP1.1, with three homoeologs in bread wheat (*Ta*MFP1.1-A1 - TraesCS1A02G119700; *Ta*MFP1.1-B1 - TraesCS1B02G139100; *Ta*MFP1.1-D1 - TraesCS1D02G120600) and two in durum wheat (*Tt*MFP1.1-A1- TRITD1Av1G054690; *Tt*MFP1.1-B1 - TRITD1Bv1G062760)(Figure S1). These had the typical three-exon structure previously reported for Arabidopsis MFP1. The other set, referred to as MFP1.2, is encoded on group 3 chromosomes, with three homoeologs in bread wheat (*Ta*MFP1.2-A1 - TraesCS3A02G117800; *Ta*MFP1.2-B1 - TraesCS3B02G137100; *Ta*MFP1.2-D1 - TraesCS3D02G119600) and two in durum wheat (*Tt*MFP1.2-A1- TRITD3Av1G038460; *Tt*MFP1.2-B1 - TRITD3Bv1G047250). MFP1.2 genes have the same three exons as MFP1.1, although there was consistently a second gene model predicted with an additional exon before the canonical second exon (Figure 1A, S1). Comparing MFP1.1 homoeologs from bread and durum wheat, the amino acid sequences shared >94% identity both within and between species (Figure S2). Similar values were obtained for MFP1.2 homoeologs. When comparing MFP1.1 to MFP1.2 homoeologs, the two paralogs shared 50-52% identity, suggesting that they are highly similar. In contrast, the wheat MFP1 sequences shared <9% identity to the Arabidopsis protein.

**Figure 1:**
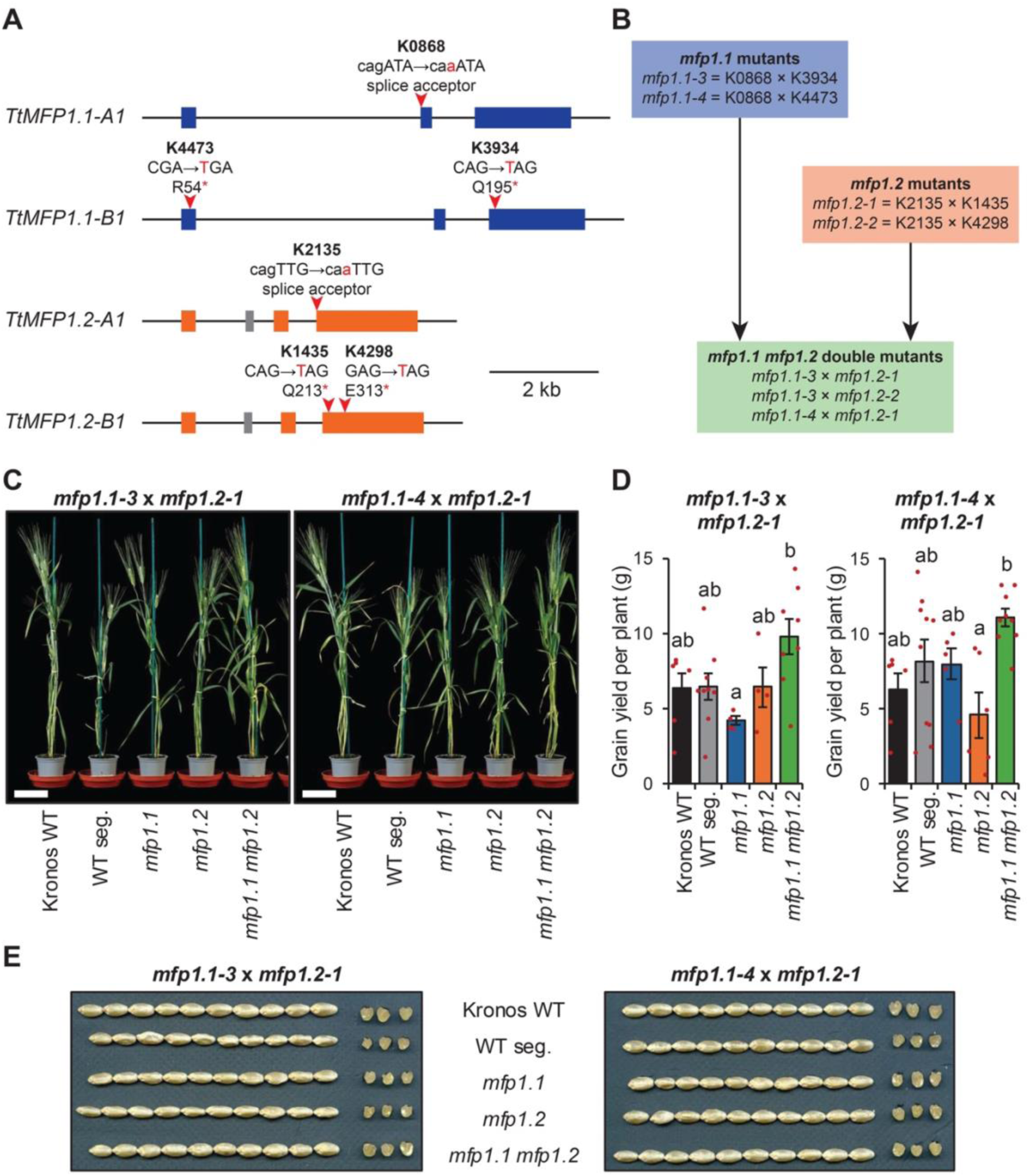
*mfp1.1 mfp1.2* mutants have normal plant growth and grain phenotypes. **A**) Gene models of *MFP1.1* and *MFP1.2* paralogs in durum wheat obtained from Svevo genome. Exons are depicted in blue (*MFP1.1*) and orange (*MFP1.2*). The putative additional exon in *MFP1.2* gene models is shown in grey. The position of the mutations in the TILLING mutants is indicated with red arrows. **B**) Different combinations of TILLING mutants used to make *mfp1.1* and *mfp1.2* mutants are shown in the blue and orange boxes, respectively. The crossing scheme for *mfp1.1* x *mfp1.2* mutants are shown in the green box. **C**) Photograph of plants at maturity. The Kronos wild type (WT) was compared with genotypes segregated from both *mfp1.1-3* x *mfp1.2-1* and *mfp1.1-4* x *mfp1.2-1* crosses: including the wild-type segregant control (WT seg.), mutants in both homoeologs of either MFP1.1 (*mfp1.1*) or MFP1.2 (*mfp1.2*), and all homoeologs of both genes (*mfp1.1 mfp1.2*). Bar = 10 cm. **D**) Grain yield per plant. Bars represent the mean ± standard error of the mean from n = 4-10 plants, while individual data points are shown in red. Values with different letters are significantly different under a one-way ANOVA and Tukey’s post hoc test at p < 0.05. **E**) Photographs of grains and transverse sections of the grain.

Using our previously generated RNAseq data for developing endosperm of durum wheat (Chen et al., 2023), we detected transcripts for both MFP1 paralogs at all timepoints between 6-30 dpa (Figure S3). For MFP1.1, transcripts peaked ∼8-10 dpa, but the A-genome homoeolog had ∼2-fold higher transcripts than the B-genome homoeolog. For MFP1.2, transcripts peaked at 10 dpa, and the expression of the two homoeologs were equal. When compared with expression patterns of known genes involved in B-type granule initiation, the expression pattern most closely followed PHS1, which also peaked at 10 dpa, while BGC1 peaked later at 18 dpa.

To allow functional characterisation of the paralogs, we used the wheat TILLING mutant collection in durum wheat cultivar Kronos (Krasileva et al., 2017) to find lines with mutations in each of the homoeologs. These mutations caused premature stop or loss of a splice acceptor site, making them likely to knock out gene function (Figure 1A). We then conducted a series of crosses to produce sets of mutants defective in both homoeologs of either MFP1.1 (referred to as *mfp1.1-3* and *mfp1.1-4*) or MFP1.2 (referred to as *mfp1.2-1* and *mfp1.2-2*). These single mutants were subsequently crossed together to create three double mutants defective in all homoeologs of both MFP1.1 and MFP1.2 (Figure 1B). We subsequently analysed in detail two sets of lines derived from the crosses: *mfp1.1-3* x *mfp1.2-1* and *mfp1.1-4* x *mfp1.2-1.* From the progeny of each of these two crosses, we selected different genotypes, including the wild-type segregant control (WT seg.), mutants in both homoeologs of either MFP1.1 (*mfp1.1*) or MFP1.2 (*mfp1.2*), and all homoeologs of both genes (*mfp1.1 mfp1.2*).

We first examined plant and grain phenotypes of the MFP1 mutants. Overall, we did not observe any significant effect of *mfp1* mutations on these phenotypes. While there was variation in plant growth among individuals, there were no differences that were consistently associated with genotype (Figure 1C). Similarly, while some significant differences in grain yield per plant were detected when comparing across genotypes, none of these were consistently observed between the *mfp1.1-3* x *mfp1.2-1* and *mfp1.1-4* x *mfp1.2-1* lines, and thus cannot be attributed to mutations in MFP1 (Figure 1D). The grains of the mutants were also visually indistinguishable from the wild type (Figure 1E), and there were no significant differences in thousand grain weight (TGW) or grain size parameters (Figure S4).

### mfp1.1 mfp1.2 mutants have altered granule size distributions

We then examined whether the mutations in *MFP1.1* and *MFP1.2* affect starch granule number and size in the endosperm. We first purified the starch from mature grains and examined starch granule size distributions using a particle size analyser (Coulter counter). While the mutation of either *MFP1.1* or *MFP1.2* homoeologs alone did not affect granule size distribution, the *mfp1.1 mfp1.2* mutants defective in both paralogs showed a distinct distribution - where the peak corresponding to B-type granules was shifted to a larger size range (Figure 2A). This shift was consistently observed in both *mfp1.1-3* x *mfp1.2-1* and *mfp1.4* x *mfp1.2-1* crosses. Curve fitting analysis of the size distributions showed that the *mfp1.1 mfp1.2* genotype, but not the single paralog mutant genotypes (*mfp1.1* or *mfp1.2*), had a significant increase in B-type granule size compared with the WT controls (Figure 2B). Interestingly, this effect was specific to the B-type granules, as there were no significant differences in A-type granule size compared with the WT controls (Figure 2C).

**Figure 2:**
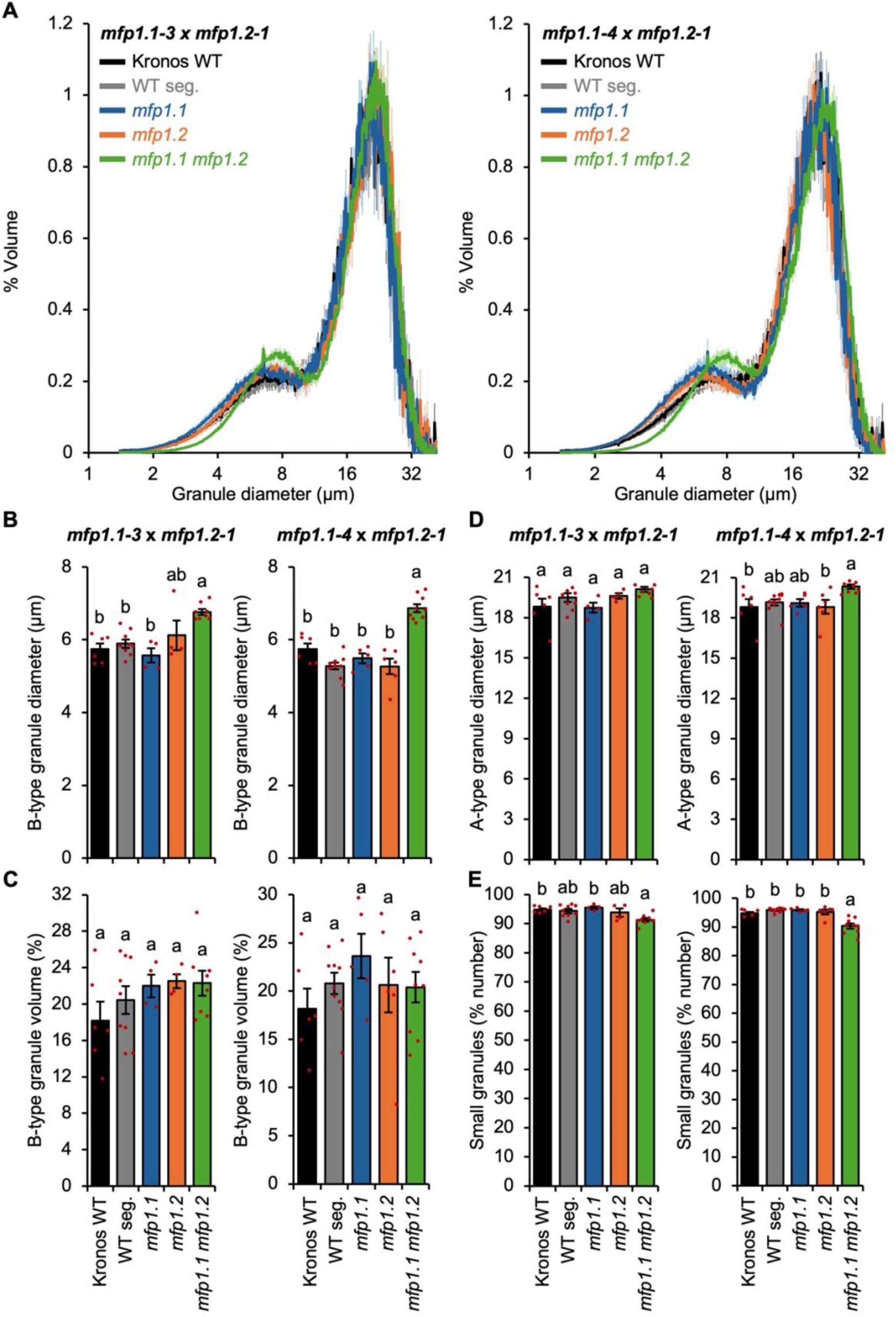
Endosperm starch from the *mfp1.1 mfp1.2* mutants have fewer but larger B-type granules. The Kronos wild type (WT) was compared with genotypes segregated from both *mfp1.1-3* x *mfp1.2-1* and *mfp1.1-4* x *mfp1.2-1* crosses: including the wild-type segregant control (WT seg.), mutants in both homoeologs of either MFP1.1 (*mfp1.1*) or MFP1.2 (*mfp1.2*), and all homoeologs of both genes (*mfp1.1 mfp1.2*). **A**) Granule size distributions were determined using a Coulter counter, and the data were expressed as relative % volume (of total starch) vs. granule diameter plots. **B**) Average diameter of B-type granules. **C**) B-type granule volume (% of total starch volume). **D**) Average diameter of A-type granules. In panels **B**-**D**, data were extracted from the relative volume vs. diameter plots in panel **A** by fitting a bimodal mixed normal distribution. **E**) The percentage of small granules by number (defined as number of granules <10 µm, as a percentage of total granule number) was calculated from the Coulter counter data. For panels A-E, plots show the mean from the analysis of n=4-10 replicate starch extractions, each using grains from a separate plant. The shading (panel A) and error bars (panels B-E) represent the standard error. Values with different letters are significantly different under a one-way ANOVA with Tukey’s post-hoc test (p < 0.05).

When B-type granule content was calculated as a percentage volume of the total starch volume, there were no significant differences between the genotypes (Figure 2D). However, since the B-type granules were on average larger in *mfp1.1 mfp1.2* mutants than the wild-type controls, we reasoned that they must be fewer of them in the mutants, such that their total volume remains comparable. Therefore, we estimated the number of B-type granules by calculating the relative number of small granules (<10 µm, which would include mostly B-type granules). This number was significantly lower in the *mfp1.1 mfp1.2* mutants when compared with the Kronos wild type in both crosses (Figure 2E), and when compared with the WT segregant in the *mfp1.1-4* x *mfp1.2-1* cross. However, in the *mfp1.1-3* x *mfp1.2-1* cross, the value was not significantly different when compared the WT segregant, likely due to large variation in the latter. These changes in granule size distribution occurred without significant differences in the total starch content of the grains (Figure S5).

Since the *mfp1.1-3* x *mfp1.2-1* and *mfp1.1-4* x *mfp1.2-1* crosses used the same mutant alleles for *mfp1.2*, we verified our findings using a third cross, *mfp1.1-3* x *mfp1.2-2*, where we isolated the WT segregant and *mfp1.1 mfp1.2* genotypes (Figure 1B). Consistent with the other crosses, we observed significantly fewer and larger B-type granules in the *mfp1.1 mfp1.2* genotype compared with the WT controls (Figure S6). Taken together, our data suggests that in wheat, loss of both MFP1 paralogs causes the formation of fewer, but larger B-type starch granules.

Next, we analysed starch granule morphology using Scanning Electron Microscopy (SEM) (Figure 3). We saw no obvious morphological differences in A- or B-type granules, suggesting that MFP1 is dispensable for granule morphogenesis. We then quantified the amylose content and amylopectin chain-length structure. We observed no differences in amylose content in any of the genotypes (Figure S5). Curiously, amylopectin chain-length distribution was altered in *mfp1.1* and *mfp1.1 mfp1.2* genotypes, with an increased proportion of chains between DP 7-11. While this effect was small, it was highly reproducible, being observed consistently in all three lines. This effect was not observed in *mfp1.2* mutants, suggesting that MFP1.1 deficiency only affects chain-length distribution.

**Figure 3:**
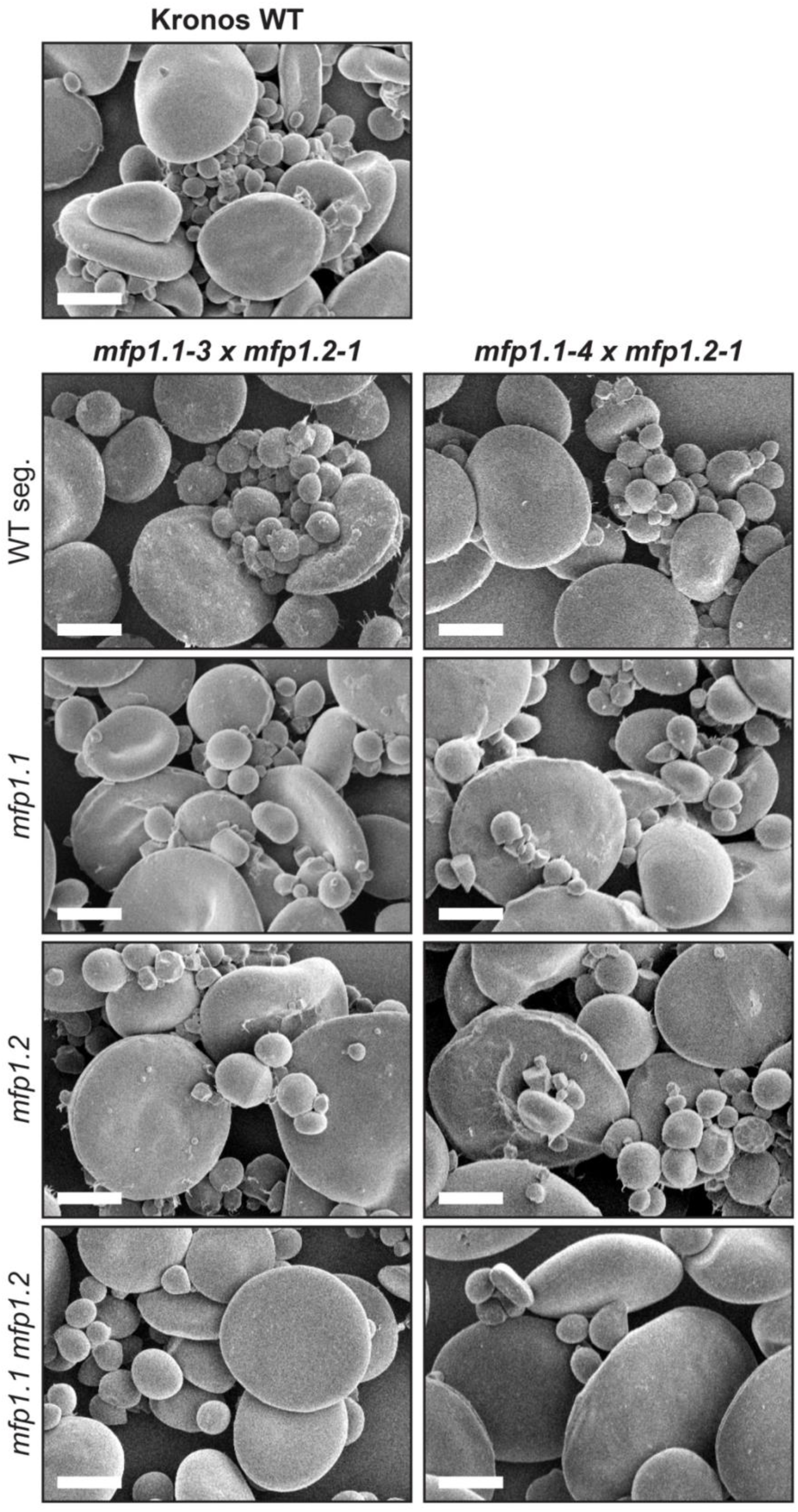
Starch granule shape is not altered in the *mfp1.1 mfp1.2* mutants. Scanning electron micrographs of purified endosperm starch. Bar = 10 µm.

Overall, our findings demonstrate that MFP1 paralogs are essential for normal starch granule size distributions in the wheat endosperm, where they primarily affect the number of B-type starch granules. Importantly, mutations in both MFP1.1 and MFP1.2 are required to observe effects on the size distribution. The fact that mutation of either paralog alone is insufficient for this starch granule phenotype suggests that the two paralogs are completely functionally redundant in their role in B-type granule number. This is in contrast to the minor role of MFP1.1 in amylopectin chain-length structure, where MFP1.2 appears not to be involved.

### Both paralogs of MFP1 interact with BGC1

Given that MFP1 is important for B-type granule initiation, we explored its interactions with BGC1 – a protein that plays a central role in granule initiation. In our previous work, we reported on a pulldown experiment where recombinant BGC1 was incubated with durum wheat endosperm extracts, and co-purifying proteins were detected using mass spectrometry (Kamble et al., 2023). There, both MFP1.1 and MFP1.2 were detected in almost equal abundance, and were the interacting proteins with the highest abundance. As this was a large-scale experiment, we tested whether both MFP1 paralogs could indeed interact with BGC1. For these experiments, we used the protein sequences from the bread wheat Chinese Spring genome. First, we used yeast-2-hybrid (Y2H), which showed positive colony growth when BGC1 was combined with either MFP1.1 or MFP1.2, suggesting direct physical interaction between the proteins (Figure 4A). We then tested these interactions using immunoprecipitation (IP) of epitope-tagged proteins transiently expressed in *Nicotiana benthamiana* (Figure 4B). In the IP using anti-YFP beads, BGC1-RFP copurified with both MFP1.1-YFP and MFP1.2-YFP, but not with free plastid-tageted YFP. Likewise, free plastidial RFP did not co-purify with MFP1.1-YFP and MFP1.2-YFP, demonstrating specific interactions between BGC1 and both MFP1 paralogs.

**Figure 4:**
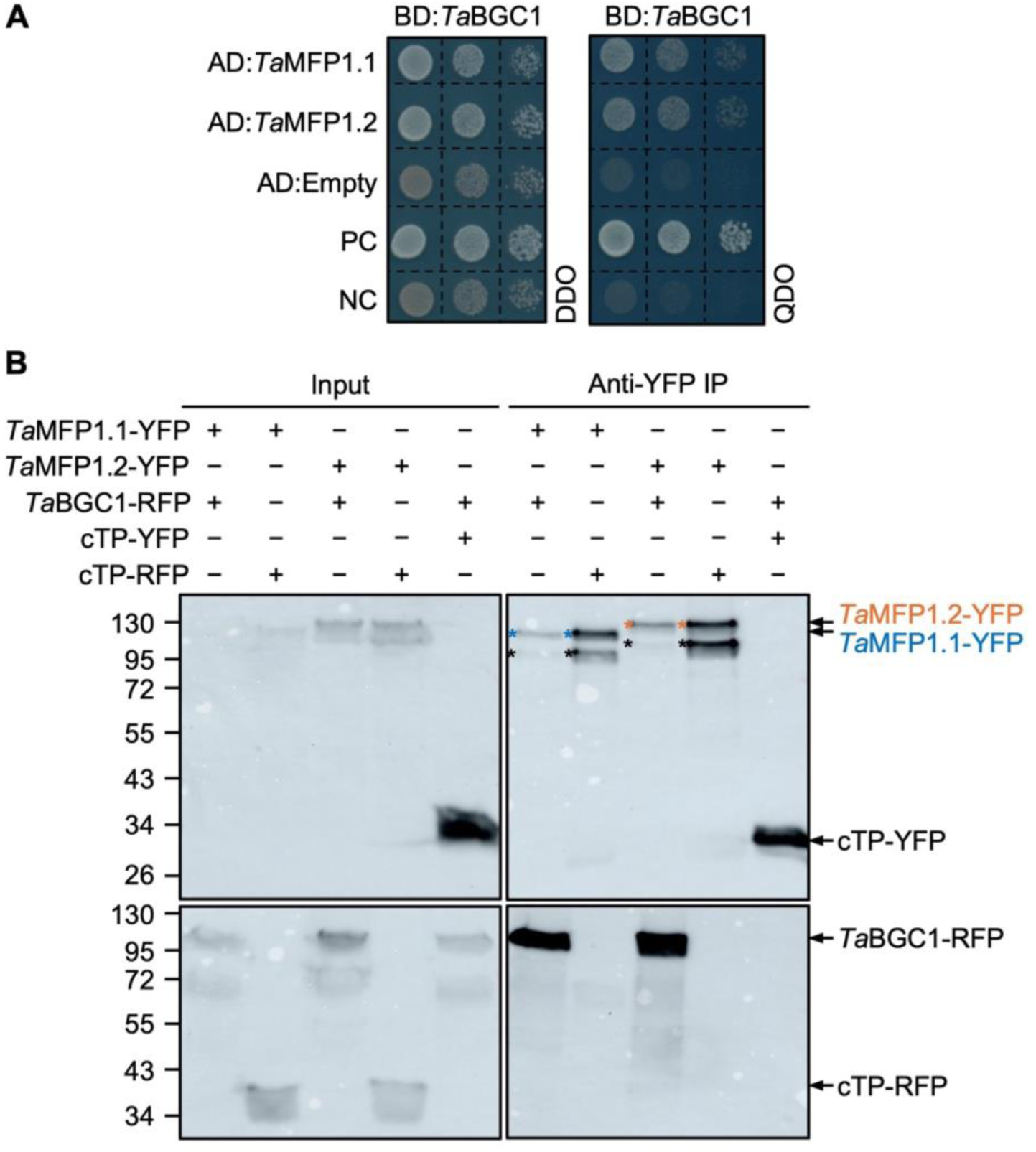
MFP1.1 and MFP1.2 physically interact with BGC1. All proteins are from the bread wheat Chinese Spring genome. **A**) Yeast two-hybrid (Y2H) assay showing interactions between AD:*Ta*MFP1.1 or AD:*Ta*MFP1.2 and BD:*Ta*BGC1, where AD is the activation domain and BD is the binding domain. Cells were plated on double dropout medium (DDO; -Leu-Trp) to analyse growth and quadruple dropout medium (QDO; -Ade-Leu-Trp-His) to analyse the interaction. PC, positive control (T-antigen and p53); NC, negative control (T-antigen and lamin). **B**) Pairwise immunoprecipitation (IP) of *Ta*MFP1.1-YFP or *Ta*MFP1.2-YFP and *Ta*BGC1-RFP co-expressed in *N. benthamiana* leaves, using anti-YFP beads. Blots of input and IP samples were probed with YFP (top panels) and RFP antibodies (bottom panels). Chloroplast-targeted YFP (cTP-YFP) and RFP (cTP-RFP) were used as controls to exclude non-specific binding to the fluorescent protein tags.

### Both wheat MFP1.1 and MFP1.2 can facilitate granule initiation in chloroplasts

Since our experiments suggested that wheat MFP1.1 and MFP1.2 were functionally redundant, we explored whether both paralogs retained the functions described for Arabidopsis MFP1. We first examined the localisation of bread wheat MFP1-YFP and BGC1-RFP proteins in *Nicotiana benthamiana* leaves. As previously reported for Arabidopsis MFP1, both wheat MFP1.1-YFP and MFP1.2-YFP had a punctate localisation pattern in *Nicotiana* chloroplasts (Figure 5A). However, we also observed ring-like structures, suggesting that the MFP1 proteins also localised to the surface of starch granules. Demonstrating a major difference from its Arabidopsis ortholog, PTST2, wheat BGC1-RFP was primarily located in ring-like structures, with only a few puncta, suggesting starch localisation. This primary starch localisation for BGC1 when expressed in *N. benthamiana* chloroplasts is consistent with our previous work (Kamble et al., 2023). The MFP1-YFP and BGC1-RFP signals in the ring-like structures showed tight co-localisation, whereas those in the puncta did not co-localise. These different localisation patterns suggests that there could be some functional divergence between the wheat and Arabidopsis proteins. However, there were no obvious differences between the localisation patterns of MFP1.1 and MFP1.2 in this heterologous system.

**Figure 5:**
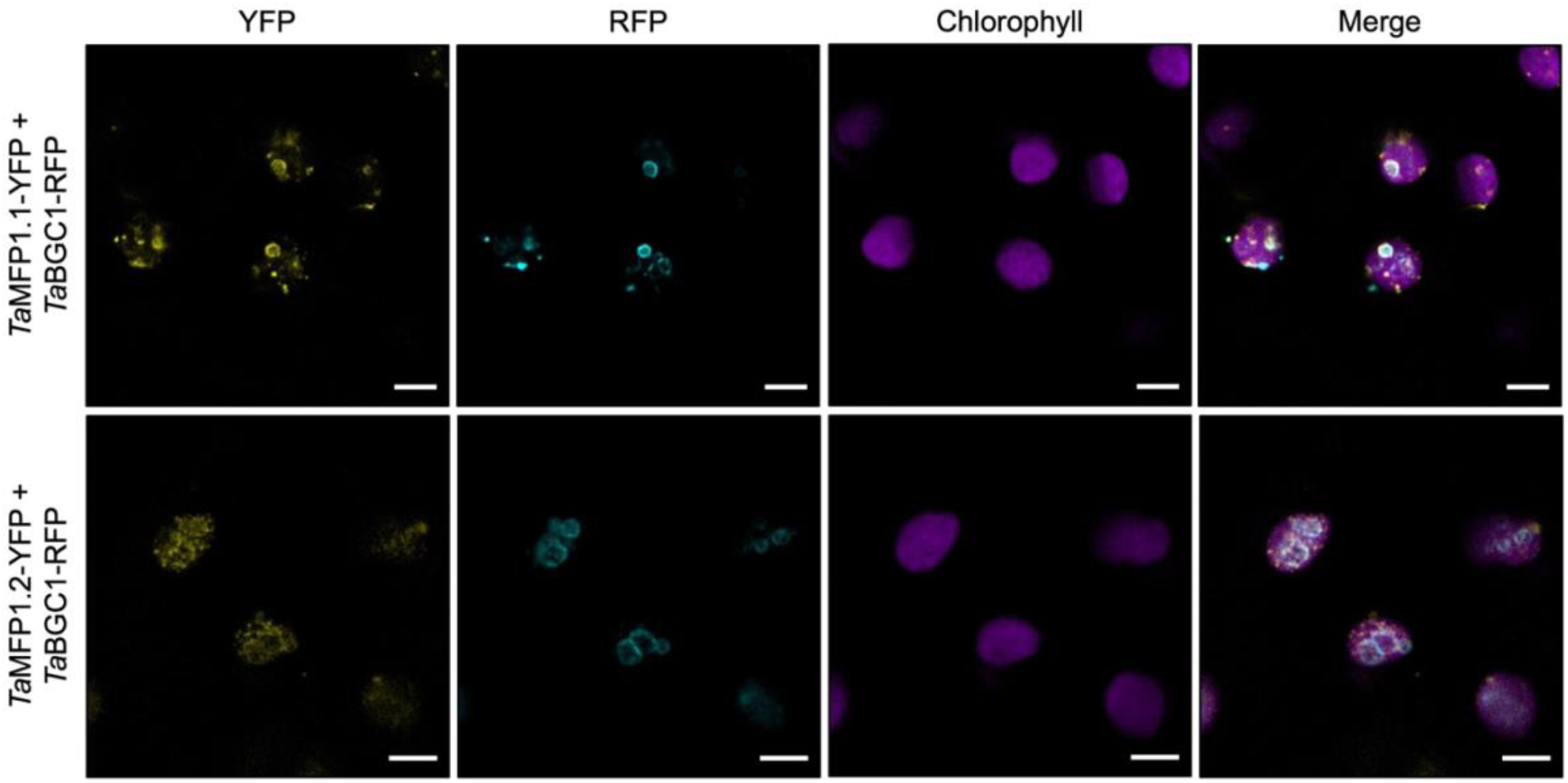
Co-localisation of wheat MFP1.1 or MFP1.2 with BGC1 in *Nicotiana benthamiana*. *Ta*MFP1.1-YFP and *Ta*MFP1.2-YFP were transiently expressed in *N. benthamiana* leaves under a Ubiquitin10 promoter, while BGC1-RFP was expressed under a 35S promoter. All proteins are from the bread wheat Chinese spring genome. Images were captured by confocal laser-scanning microscopy. Chlorophyll autofluorescence was used to locate the chloroplasts. Bars = 5 μm.

We then made transgenic Arabidopsis lines expressing either wheat *Ta*MFP1.1 or *Ta*MFP1.2 in the *mfp1* mutant background. Both wheat MFP1 paralogs were expressed under the constitutive Ubiquitin10 promoter, and with a C-terminal YFP tag. We screened the lines for transgene expression using antibodies against MFP1.1 and MFP1.2. The fusion proteins were visible around their expected sizes (MFP1.1-YFP: 113.1 kDa; MFP1.2-YFP: 116.8 kDa)(Figure 6A). We used light microscopy to examine the granule number per chloroplast. While the *mfp1* mutant had mostly single starch granules per chloroplast, the expression of either wheat MFP1 isoform could restore the synthesis of multiple granules per chloroplast, but not to the same number seen in the WT. These results suggest that both wheat MFP1 paralogs can promote granule initiation in Arabidopsis chloroplasts, but they do so with low efficiency compared with the Arabidopsis MFP1.

**Figure 6:**
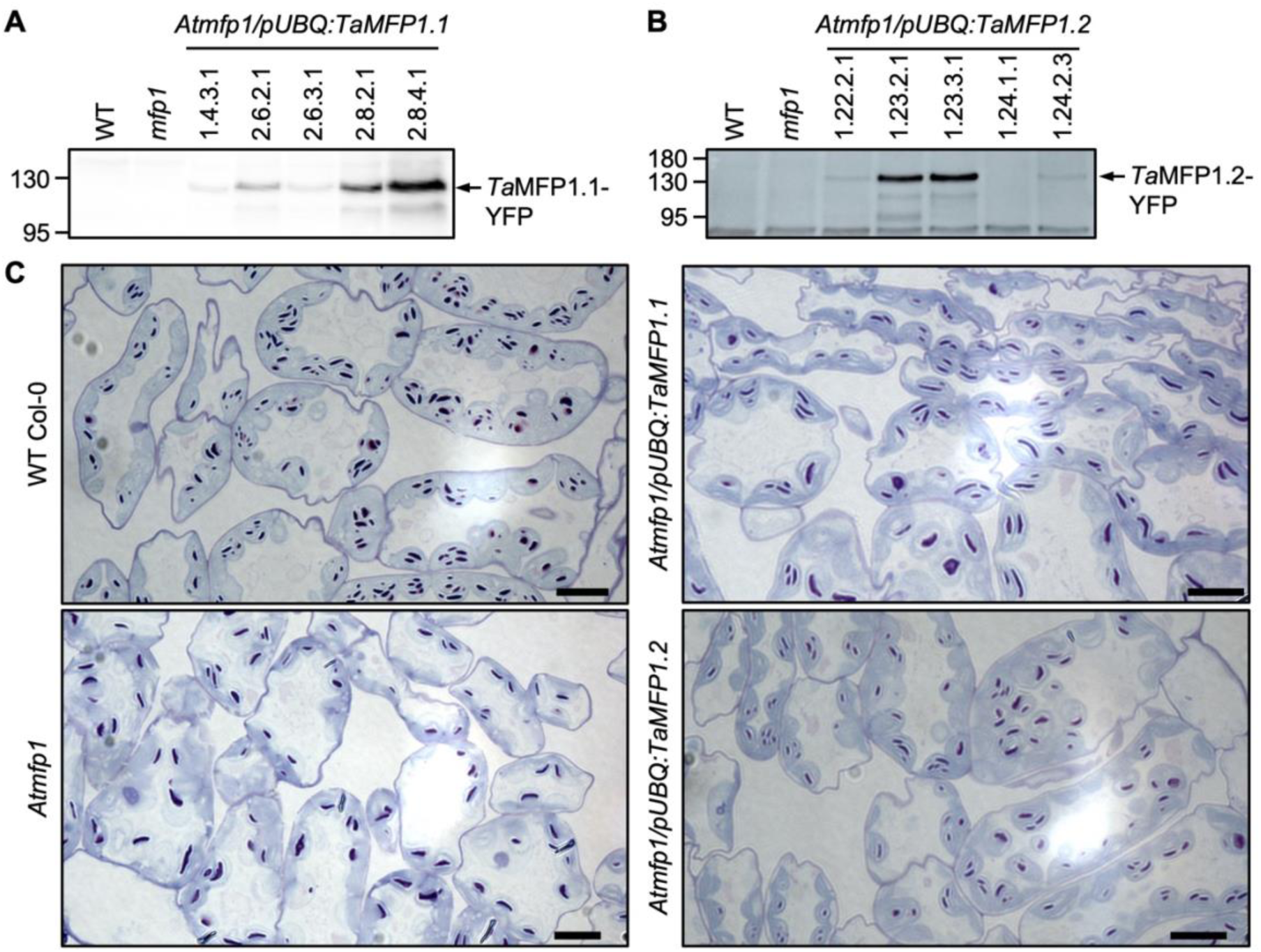
Wheat MFP1.1 and MFP1.2 can partially complement starch granule morphology phenotypes of the Arabidopsis *mfp1* mutant. **A**) Immunoblots from independent transgenic Arabidopsis lines expressing wheat *Ta*MFP1.1-YFP or *Ta*MFP1.2-YFP under the Ubiquitin10 promoter (pUBQ) in the *Atmfp1* mutant background (*Atmfp1*/pUBQ:*Ta*MFP1.1 or *Atmfp1*/pUBQ:*Ta*MFP1.2). Total proteins were extracted from leaves and immunoblotted using anti-*Ta*MFP1.1 or anti-*Ta*MFP1.2 antibodies. Lanes were loaded on an equal protein basis (40 µg). The migration of molecular weight markers are indicated in kilodaltons (kDa) to the left of each panel. **B**) Starch granules in chloroplasts observed using light microscopy for the selected lines analyzed in (A) of the wild-type (Col-0), *Atmfp1* mutant, and independent transgenic lines expressing wheat *Ta*MFP1.1-YFP or *Ta*MFP1.2-YFP in the *Atmfp1* background. Thin sections were stained with toluidine blue and periodic acid/Schiff’s reagent. Bars = 10 µm.

## Discussion

### MFP1 is required for normal B-type starch granule initiation

We found that MFP1 is required for normal B-type granule initiation in wheat. Mutants deficient in both MFP1.1 and MFP1.2 had fewer, but larger B-type granules (Figures 2, S6). The loss of both MFP1 paralogs did not affect either the total starch content of the grain or grain size (Figures S4, S5). Since fewer B-type granules were initiated in the mutant grains, it is possible that each B-type granule was able to grow larger because there were fewer granules competing for substrates. Such a trade-off between B-type granule number and size has been observed previously in the *mrc* K2485 mutant, which also has reduced frequency of B-type granule initiations (Fahy et al., 2025). Curiously, no change in A-type granule size was detected in the *mfp1.1 mfp1.2* mutants, even though having fewer B-type granules would also likely provide more substrates for A-type granule growth. However, this may only result in a small change in A-type granule size, which may not be detectable. Even the *bgc1-1* mutant with almost no B-type granules has no detectable change in A-type granule size (McNelly et al., 2025). Both A- and B-type granules retained their typical lenticular and round morphologies, respectively (Figure 3). This suggests that the role of MFP1 in granule initiation appears to exclusively affect B-type granules.

It is likely that B-type granule initiation is reduced in the mutant is due to the loss of MFP1 affecting BGC1 function. We demonstrated that both MFP1.1 and MFP1.2 can interact with BGC1 (Figure 4), suggesting that the MFP1-PTST2 interaction reported in Arabidopsis is conserved in both wheat MFP1 paralogs (Seung et al., 2018). In wheat, BGC1 plays an important role in both A- and B-type granule initiation, but B-type granule initiation is particularly sensitive to changes in BGC1 function, allowing a range of mutant phenotypes depending on the degree of BGC1 inactivation. Complete loss of BGC1 causes strong defects in A-type granule formation, due to supernumerary initiations during A-type granule initiation, and almost no normal B-type granule initiation (Chia et al., 2020; Hawkins et al., 2021). However, mutants with reduced BGC1 function can have normal A-type granule formation with severely reduced numbers of B-type granules (Chia et al., 2020; McNelly et al., 2025). The reduced BGC1 function can be achieved through reductions in gene dosage (i.e., mutating most but not all homoeologs), or through hypomorphic mutations. For example, the *bgc1-1* mutant of Kronos, which has a premature stop codon mutation in the A-genome copy of *BGC1* and a V470I amino acid substitution in the B-genome copy, produces almost no B-type granules (Chia et al., 2020). Therefore, it is plausible that the loss of MFP1 paralogs reduces, but does not abolish BGC1 function, leading to the observed reduction in B-type granule number in *mfp1.1 mfp1.2* mutants. However, it must be noted that the overall phenotype of the *mfp1.1 mfp1.2* mutants is milder than *bgc1-1*, as they still produce many B-type granules. MFP1 therefore must play a role that promotes the formation of normal numbers of B-type granules, but is not strictly essential for the initiation process.

### Shared and diverged functions between MFP1 paralogs in granule initiation and amylopectin structure

MFP1.1 and MFP1.2 arose from a gene duplication in the grasses (Figure S1), providing opportunity for functional divergence between paralogs. However, we found no signs of functional divergence in their role in B-type granule initiation, where both paralogs appear completely redundant. Both MFP1.1 and MFP1.2 are expressed in the endosperm with comparable transcript levels (Figure S3). Both paralogs can also interact with BGC1. Mutating either paralog alone had no effect on B-type granule initiation due to compensation from the other paralog, since the effect on B-type granule number and size was only observed when both paralogs were mutated.

Interestingly, we discovered an unexpected role of MFP1 in amylopectin chain length distribution where only MFP1.1 contributes, demonstrating some functional divergence between the two paralogs. We observed a small enrichment in chains DP 7-11 in all *mfp1.1* and *mfp1.1 mfp1.2* genotypes. While the change is small, suggesting only a minor role, it was consistently observed in all three lines. The *mfp1.2* mutation did not alter chain length distribution, neither by itself nor in the *mfp1.1* mutant backgrounds, suggesting that it does not contribute at all to the process. The effect on chain length distribution is also therefore unlikely to be a consequence of altered granule initiation, since the *mfp1.1* mutants had altered chain length distribution in the absence of detectable changes in B-type granule size/number.

Currently, it is not possible to say whether MFP1.1 gained a role in determining amylopectin structure in wheat, or whether MFP1.2 lost the role. However, this could be determined by investigating whether the effect of MFP1 on chain length distribution is also observed in other species, for instance by analysing the Arabidopsis *mfp1* mutant. Also, how MFP1, a non-catalytic protein that primarily consists of a coiled coil, influences amylopectin structure remains to be determined. Since some starch-bound MFP1 was detected after transient expression in *N. benthamiana* (Figure 5), it is plausible that it indirectly affects amylopectin structure by influencing the activity of other biosynthetic enzymes, for example Starch Synthase 1 which influences the proportion of short amylopectin chains (Mcmaugh et al., 2014; Trafford et al., 2026).

### MFP1 requirement in non-photosynthetic amyloplasts

Our work demonstrates a role for MFP1 in starch granule initiation in non-photosynthetic amyloplasts. This is important considering that the role of MFP1 in starch synthesis was originally established in Arabidopsis chloroplasts, which has a vastly different organelle structure and granule initiation pattern compared with wheat amyloplasts. In chloroplasts, granules form within stromal pockets in between thylakoid membranes (Burgy et al., 2021; Esch et al., 2022; Ilse et al., 2025). The pockets represent defined locations where granules initiate. It is proposed that MFP1 defines the locations of granule initiations by controlling PTST2 localisation (Seung et al., 2018; Sharma et al., 2024).

Central to its role in defining initiation locations is the localisation of MFP1, since altering MFP1 localisation by targeting it to other areas of the chloroplast can also alter the location of starch granules (Sharma et al., 2024). MFP1 is bound to the stromal side of thylakoid membranes, and in Arabidopsis it is exclusively found in membrane-bound protein fractions (Jeong et al., 2003; Seung et al., 2018). Thus, PTST2 in Arabidopsis is also partially bound to thylakoids through its interaction with MFP1 (Seung et al., 2018). However, in wheat amyloplasts, there are no internal thylakoid membranes, and thus a role for MFP1 in defining specific regions for granule initiation is less relevant, especially for A-type granules which form from a single granule initiation per amyloplast during early grain development. Consistent with the diminished importance of MFP1 in amyloplasts, the phenotype of the wheat *mfp1.1 mfp1.2* mutants on B-type granule number is notably weaker than *bgc1* mutants. This in contrast to Arabidopsis, where the reductions in granule number per chloroplast in *mfp1* is of similar severity to *ptst2* mutants (Seung et al., 2018).

While our work clearly shows that our MFP1 mutants have a B-type granule initiation phenotype, and confirms that wheat MFP1s interact with BGC1, we are still investigating several possibilities to further elucidate the mechanism of MFP1 function in the endosperm. Firstly, we are exploring MFP1 localisation in amyloplasts, to determine whether or not MFP1 is stromal or membrane-bound in the absence of thylakoids. This is especially important since our experiments in *N. benthamina* leaves show that MFP1 can co-localise with BGC1 on starch granules, suggesting it could exist in a non-membrane-bound form (Figure 5). Secondly, it is possible that the reason why MFP1 is more important for the initiation of B-type granules than A-type granules is because at least some B-type granules are initiated in amyloplast stromules, which may require more coordination with amyloplast structure (Parker, 1985; Langeveld et al., 2000). Thus, we are exploring stromule formation in the *mfp1.1 mfp1.2* mutants. Finally, we will investigate whether the *mfp1.1 mfp1.2* mutants have defects in starch initiation in leaves. That will inform the extent to which MFP1 functions have diverged at a species-level, as opposed to at the tissue level. This is particularly pertinent since the expression of either wheat MFP1 could only partially complement the Arabidopsis *mfp1* mutant (Figure 6), and since the wheat proteins had very low amino acid identity compared with the Arabidopsis protein (Figure S2).

## Materials and Methods

### Plant materials and growth conditions

Wheat and *N. benthamiana* plants were grown in climate-controlled glasshouses at the John Innes Centre, Norwich, UK. For wheat, the glasshouses were set to provide a minimum 16 h of light at 20°C and 16°C during the dark. For *N. benthamiana*, glasshouses set to provide a minimum of 16 h light (200 μmol photons m^−2^ s^−1^) and a constant temperature (22°C) with 60% relative humidity. Arabidopsis plants were grown in controlled environment chambers (MLR-352H, PHCbi) set to a photoperiod of 16 h light (150 μmol photons m^−2^ s^−1^) and 8 h dark, constant temperature (20°C) with 60% relative humidity.

Mutant lines of MFP1.1 and MFP1.2 in durum wheat (*Triticum turgidum* cv. Kronos) from the wheat in silico TILLING resource were obtained from the John Innes Centre Germplasm Resources Unit. The lines and their mutant homoeologs are described in Supplemental Table 1. Plants were crossed to combine mutations in the A- and B-homoeologs, and lines carrying mutations in both homoeologs of MFP1.1 (*mfp1.1-3* and *mfp1.1-4*) or MFP1.2 (*mfp1.2-1* and *mfp1.2-2*) were selected in the F_2_ generation using KASP V4.0 genotyping (LGC, Teddington). The KASP primers are in Supplemental Table 1. These lines were subsequently crossed, and in the F2 generation, we isolated the *mfp1.1 mfp1.2* lines defective in all homoeologs of both paralogs (i.e., the quadruple mutant), as well as the sibling controls that were WT, or defective in both homoeologs of either MFP1.1 (*mfp1.1*) or MFP1.2 (*mfp1.2*).

### Phylogenetic analyses

For constructing phylogenetic trees, we used BLASTp against the bread wheat genome (IWGSC)(Appels et al., 2018) and the durum wheat genome (Svevo 1.1)(Maccaferri et al., 2019) to obtain sequences of MFP1 homologs. These sequences were added to the alignment in Seung et al. (2018) using MAFFT alignment (Katoh and Standley, 2013)(Supplemental File 1). The alignment was used to build a maximum likelihood tree using raxML (Edler et al., 2020), using the HIVB+G4+F model and with the *Physcomitrella* sequence being used as the outgroup, and with 1000 bootstrap replicates.

### Grain morphometrics

Grain yield per plant was quantified as the total weight of grains harvested from each plant. The thousand grain weight and grain size parameters were quantified using the MARViN seed analyser (Marvitech GmbH, Wittenburg).

### Starch purification and determination of granule morphology, size distribution

Starch purification from mature grains (2-3 grains per extraction) was performed using filtration and Percoll gradient centrifugation, exactly as described in Kamble et al. (2023).

To quantify starch granule size distributions, purified starch was suspended in Isoton II electrolyte solution (Beckman Coulter) and analysed using a Multisizer 4e Coulter counter fitted with a 70 µm aperture tube (Beckman Coulter); measuring a minimum of 100,000 particles per sample. These data were used to produce relative volume vs. diameter plots. To derive mean diameters of A- and B-type granules, and the B-type granule volume percentage (defined as volume occupied by B-type granules as a percentage of the total volume of starch), we fitted mixed bimodal curves to the relative volume vs. diameter plots using the Python analysis script described by McNelly et al. (2025), using the normal-normal curve setting. The percentage number of small granules were calculated directly from the Coulter counter data (without curve fitting), as the number of particles less than 10 µm relative to the total number of particles.

The morphology of purified granules was examined using a NanoSEM 450 (FEI) scanning electron microscope (SEM), as described in Kamble et al. (2023).

### Total starch and amylose content, and amylopectin chain length

Starch content of mature grains (in glucose equivalents) was quantified using flour (5-10 mg), by digesting starch with thermostable α-amylase and amyloglucosidase and measuring glucose using the hexokinase/glucose-6-phosphate dehydrogenase assay. This was carried out exactly as described in Hawkins et al. (2021).

Amylose content was determined on purified starch granules using iodine colourimetry, exactly as described in Kamble et al. (2023)

For amylopectin chain length distribution analysis, an equivalent of 0.1 mg starch was gelatinised and debranched with isoamylase and pullulanase, and the linear chains were analysed by High Performance Anion Exchange Chromatography with Pulsed Amperometric Detection on a Dionex ICS-5000-PAD fitted with a PA-200 column (Thermo). The exact procedures are described in Watson-Lazowski et al. (2026).

### Plasmid construction

For plasmid construction, we synthesised the coding sequences of *Ta*MFP1.1-A1 and *Ta*MFP1.2-A1 as gBlocks gene fragments (IDT) flanked with attB1 and attB2 Gateway recombination sites, using codon optimisation on the coding sequence to ease sequence complexity (Supplemental File 2). The fragments were recombined into the Gateway Entry vector pDONR221 using the BP Clonase II (Invitrogen). This yielded the entry vectors *Ta*MFP1.1:pDONR221 and *Ta*MFP1.2:pDONR221, alongside our existing *Ta*BGC1:pDONR221 (Hawkins et al., 2021) and RbcS transit peptide:pDONR221(for targeting free YFP or RFP to chloroplasts) (Kamble et al., 2023). For Y2H, we recombined the coding sequences from these entry vectors into the destination vectors pDEST-GADT7 (activation domain, AD) and pDEST-GBKT7 (binding domain, BD) (Clontech). For plant expression, we recombined the coding sequences from these entry vectors into the destination vectors pUBC-YFP (Ubiquitin10 promoter; C-terminal YFP) or pB7RWG2 (35S promoter; C-terminal RFP).

### Protein interaction analyses and Nicotiana benthamiana protein expression

For Y2H, constructs were transformed into the Y2H Gold yeast strain (Clontech) using the EZ-Yeast Transformation Kit (MP Biomedicals) according to the manufacturer’s protocol. Transformants were selected on synthetic double dropout medium (SD/−Leu/−Trp). Positive transformants were spotted onto quadruple dropout medium (SD/−Leu/−Trp/−His/-Ade) for interaction assessment. The AD-T antigen and BD-p53 combination were used as a positive control, and AD-p53 with BD-Lam was used as a negative control.

For pairwise immunoprecipitations and protein localisation, proteins were transiently expressed in *N. benthamiana* leaves through infiltration of *Agrobacterium tumefaciens* cells harbouring the relevant constructs, as previously described (Kamble and Majee, 2022). Infiltrated plants were incubated overnight in the dark followed by 2 days in a 16 h/8 h light–dark cycle. Immunoprecipitations were conducted exactly as described in Kamble et al. (2023), and analysed with immunoblotting using anti-YFP (Torrey Pines; TP401 - 1:5000) and anti-RFP (Abcam plc; ab34771 - 1:2000) primary antibodies, and horseradish peroxidase-coupled anti-rabbit secondary antibody (Sigma; A0545 - 1:15,000). For localisation, samples were imaged using a Zeiss LSM 780 Confocal Microscope with a 40X water immersion lens. YFP was excited using an argon laser at 514 nm, and emitted light was captured between 518-557 nm. RFP was excited using an argon laser at 488 nm, and emitted light was captured between 520-600 nm. Chlorophyll autofluorescence was excited with an argon laser at 514 nm, and emitted light was captured between 662-721 nm.

### Production of MFP1.1 or MFP1.2 antibodies

Antibodies specific to MFP1.1 and MFP1.2 were produced at Eurogentec using synthetic peptides, which were produced individually with a cysteine at the N-terminal to allow conjugation to Keyhole Limpet Hemocyanin (KLH). The sequences for for MFP1.1: PPRSAKKIYRRRKDR, LSSSLSTKEGDYQSL and for MFP1.2: TSQTPNEQSVNDGTQ, VDEPQTGGTQSETPL. Both peptides for each protein were co-immunised into rabbits using the Double X Speedy program. Antibodies were enriched from antiserum using protein A-agarose (Sigma-Aldrich).

### Complementation of Atmfp1 mutants

The constructs carrying YFP-tagged MFP1.1 or MFP1.2 under the Ubiquitin10 promoter was transformed into Arabidopsis by floral dipping. Transformants were selected in the T_1_ generation using the Basta resistance marker. Basta-resistant individuals from the T_2_ or T_3_ generation (heterozygous or homozygous for the transgene; single or multiple insertions) with MFP1.1 or MFP1.2 expression were confirmed using immunoblots.

For immunoblotting to check for protein expression, fully developed leaves from 4-week old Arabidopsis plants were homogenized in extraction buffer (40 mM Tris-HCl, pH 6.8, 5 mM MgCl_2_, 2% (w/v) SDS, protease inhibitor cocktail (Roche)). The homogenate was heated at 95°C for 10 min, and insoluble material was removed by centrifugation at 20,000*g* for 10 min. The concentration of proteins was determined using the BCA assay (Thermo Fisher Scientific). The following dilutions of primary antibodies were used for immunoblotting: anti-MFP1.1 - 1:2500, anti-MFP1.2 - 1:2500. Proteins were detected using chemiluminescence from horseradish peroxidase-coupled secondary antibodies Anti-rabbit HRP (Sigma; A0545 - 1:15,000).

Starch granules were visualised in leaf chloroplasts using light microscopy on semi-thin sections that were stained with toluidine blue and Periodic Acid-Schiff staining, exactly as described in Watson-Lazowski et al. (2022).

### Statistical analysis

Statistical analyses were carried out using the SPSS program (SPSS Statistics, IBM).

## Accession numbers

The accession numbers corresponding to the genes investigated in this study are:

TraesCS1A02G119700 (*TaMFP1.1-A1*), TraesCS1B02G139100 (*TaMFP1.1-B*), TraesCS1D02G120600 (*TaMFP1.1-D1*), TRITD1Av1G054690 (*TtMFP1.1-A1*), TRITD1Bv1G062760 (*TtMFP1.1-B1*), TraesCS3A02G117800 (*TaMFP1.2-A1*), TraesCS3B02G137100 (*TaMFP1.2-B1*), TraesCS3D02G119600 (*TaMFP1.2-D1*), TRITD3Av1G038460 (*TtMFP1.2-A1*), TRITD3Bv1G047250 (*TtMFP1.2-B1*), TraesCS4A02G284000 (*TaBGC1-A1*), TraesCS4B02G029700 (*TaBGC1-B1*), TraesCS4D02G026700 (*TaBGC1-D1*), TRITD4Av1G198830 (*TtBGC1-A1*), TRITD0Uv1G034540 (*TtBGC1-B1*).

## Acknowledgements

We thank JIC Horticultural Services for providing growth facilities and maintenance of plant material, JIC Bioimaging for providing access to microscopes, JIC Genotyping for providing support for KASP genotyping and JIC Chemistry for providing access to HPAEC-PAD instruments. This work was funded through a John Innes Foundation (JIF) Chris J. Leaver Fellowship (to D.S) and Biotechnology and Biological Sciences Research Council (BBSRC, UK) research grants BB/W015935/1 and BB/W01632X/2 (to D.S and K.T), as well as BB/Z517501/1 and UKRI1923 (to D.S), and BBSRC Institute Strategic Programme grants BB/X01097X/1 and BB/X011003/1 (to the John Innes Centre).

## Author contributions

N.U.K. and D.S. conceived and designed the study. All authors performed the research and analysed data. N.U.K and D.S. wrote the article with input from all authors.

## Supplemental Figures and Tables

Supplemental File 1: Sequences alignment using MAFFT alignment for constructing phylogenetic tree.

Supplemental File 2: Codon optimised coding sequences of *Ta*MFP1.1-A1 and *Ta*MFP1.2-A1.

## Supplemental Figures

**Supplemental Figure S1.**
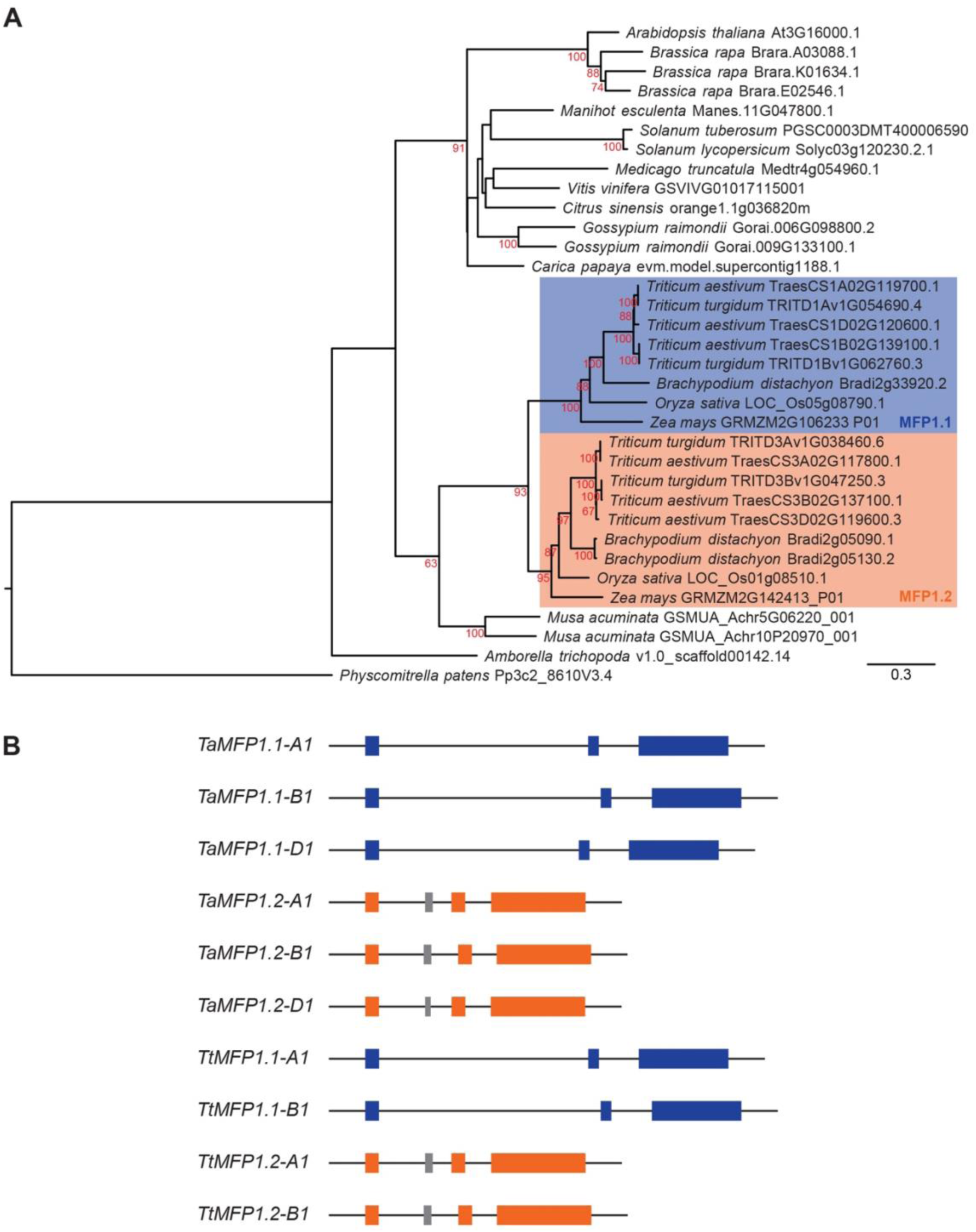
MFP1.1 and MFP1.2 result from gene duplication in the grasses. **A**) A maximum likelihood phylogenetic tree was produced using RaxML from MFP1.1 and MFP1.2 *s*equences with 1000 bootstrap replicates. Bootstrap values >50 are shown next to the nodes. The MFP1.1 clade is shaded in blue, whilst the MFP1.2 clade is shaded in orange. Branch lengths represent the number of substitutions per site, indicated by the scale bar. **B**) Gene models of MFP1.1 and MFP1.2 paralogs in bread wheat (*Triticum aestivum - Ta*) and in durum wheat (*Triticum turgidum - Tt*). Exons are depicted in blue (MFP1.1) and orange (MFP1.2). The putative additional exon in *MFP1.2* gene models is shown in grey.

**Supplemental Figure S2.**
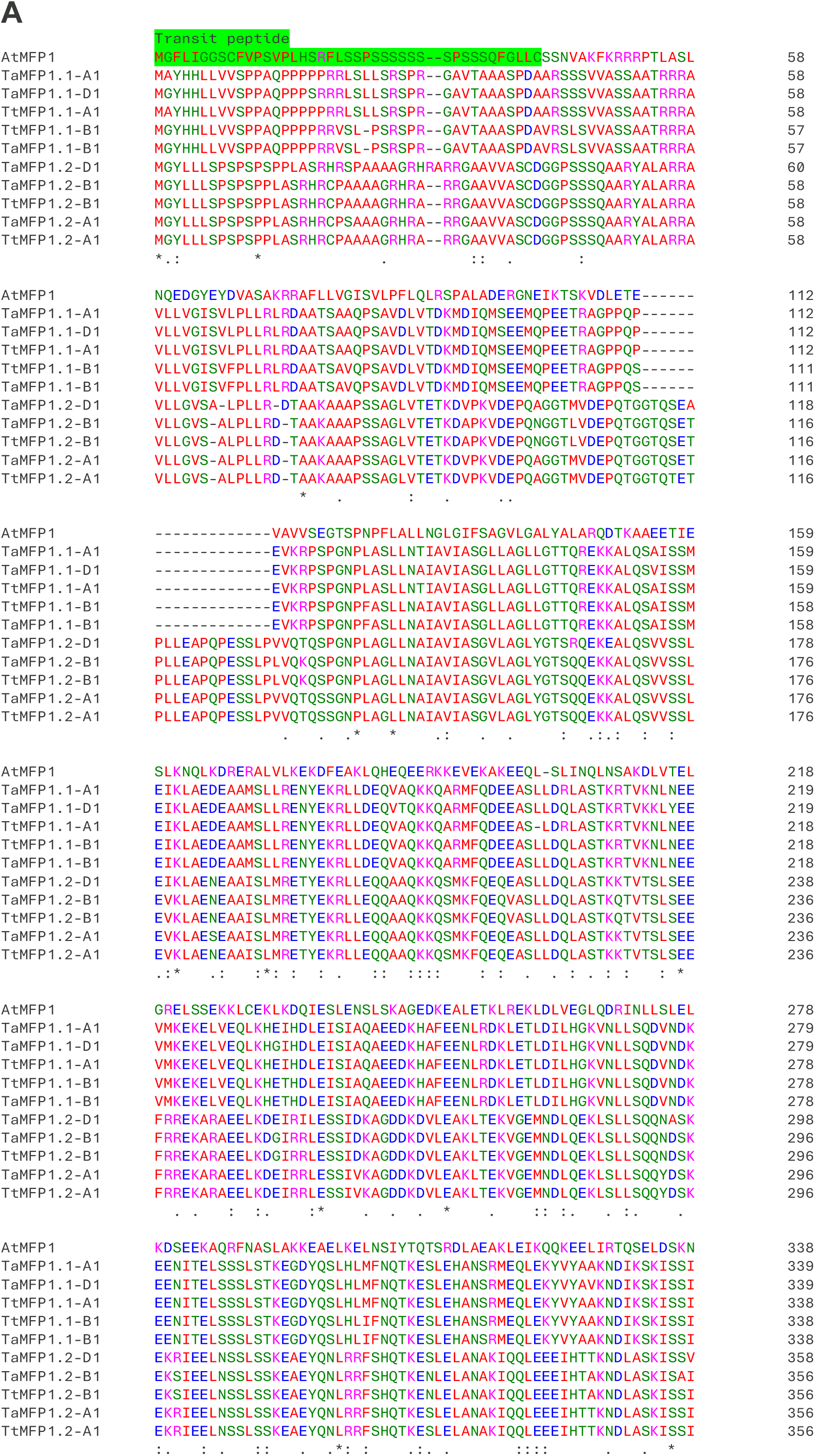
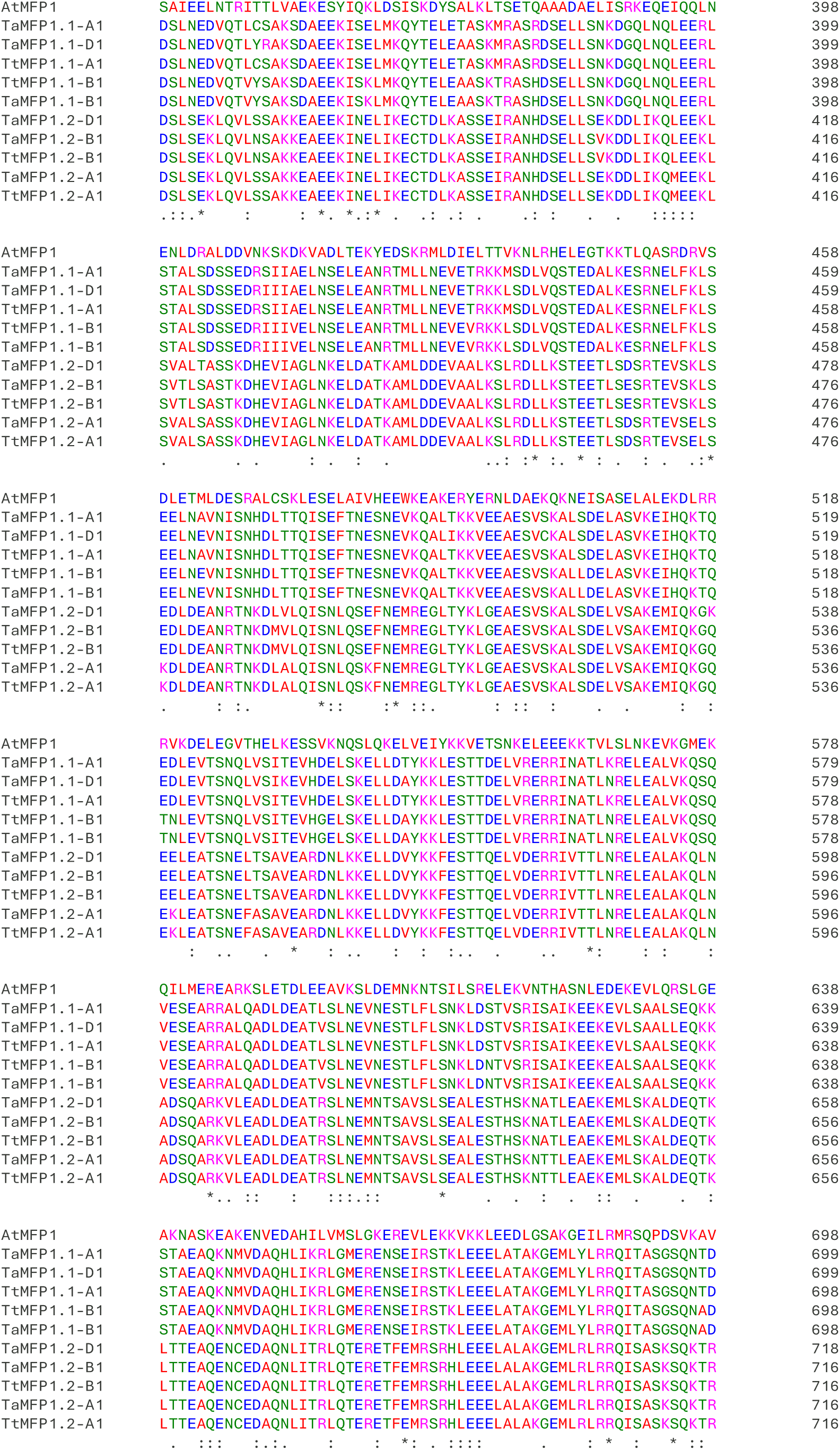
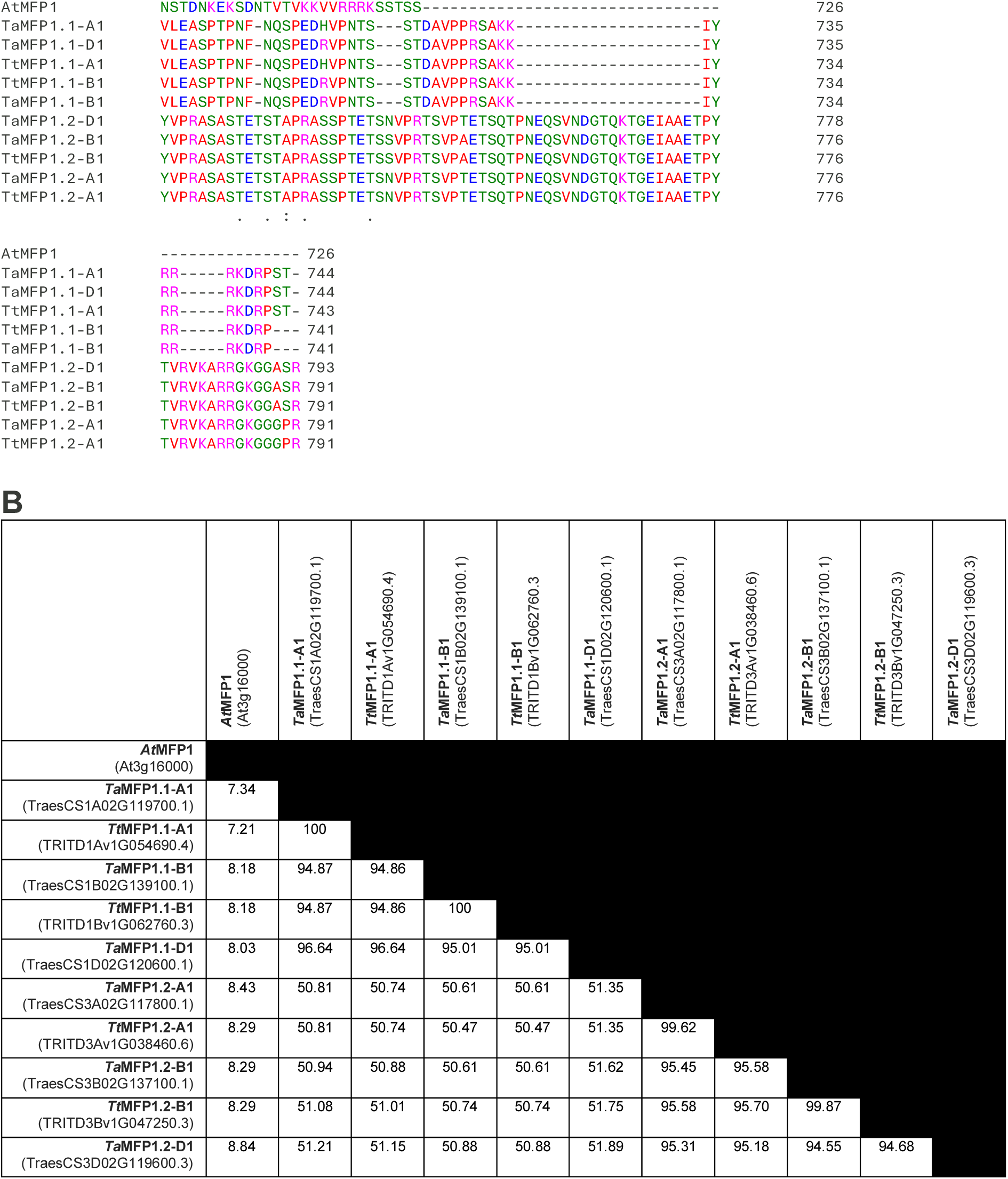
Protein sequence alignment for MFP1.1 and MFP1.2. **A)** Sequences were aligned using Clustal Omega. Region representing the chloroplast transit peptide in the Arabidopsis sequence is marked in green. Symbols under the alignment indicate full (*), strong (:) or weak (.) conservation. **B**) Matrix of percentage amino acid identity values calculated on the Clustal Omega alignment in A. *Ta* = *Triticum aestivum*, *Tt* = *Triticum turgidum*

**Supplemental Figure S3.**
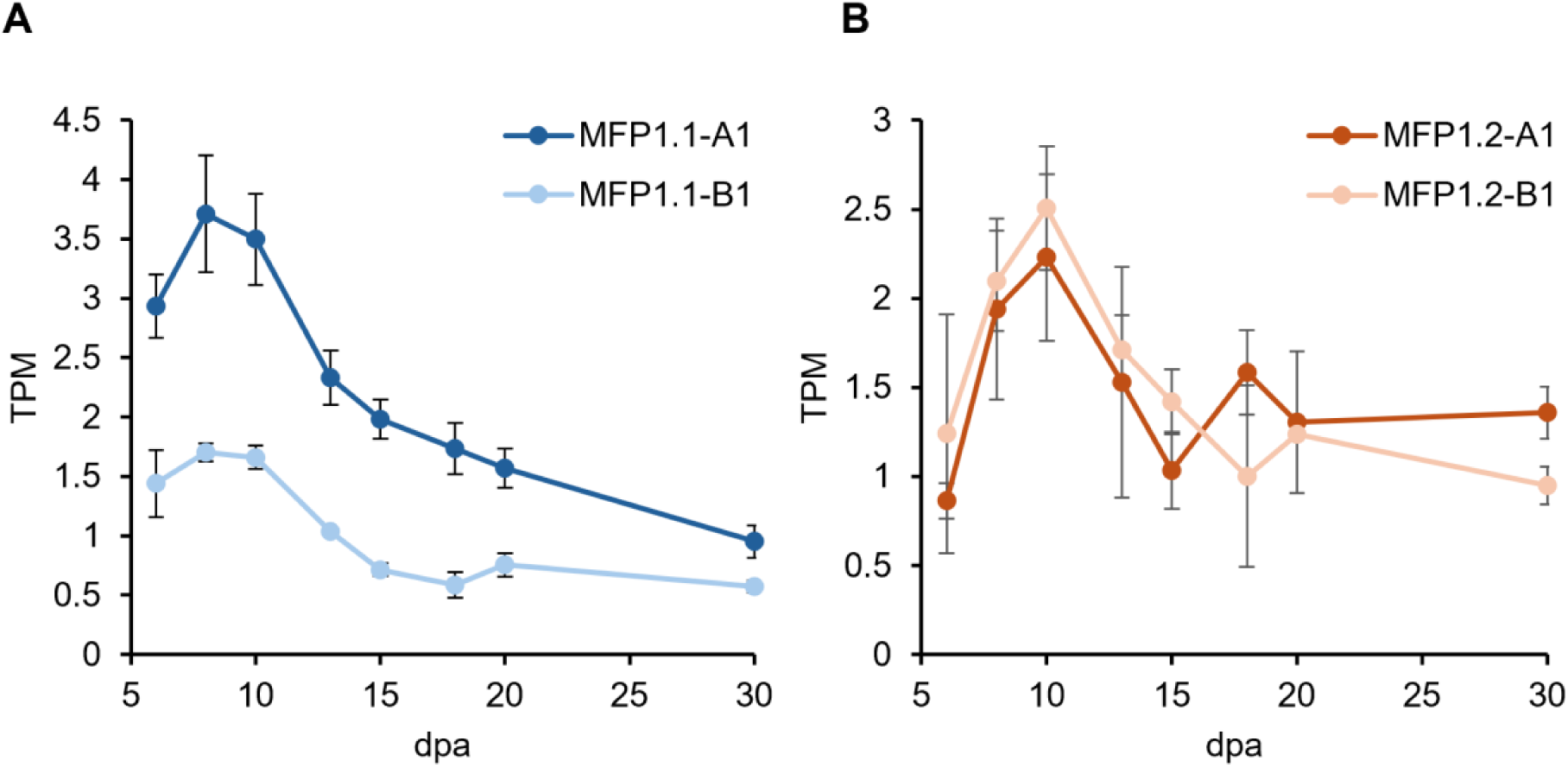
MFP1.1 and MFP1.2 expression in developing wheat grains. Expression levels of MFP1.1 and MFP1.2 paralogs in the endosperm of durum wheat across different stages of wheat grain development days post anthesis (dpa). Data are from the RNAseq study by Chen et al. (2023). Values are in transcripts per million (TPM) and are mean ± standard error of the mean from n = 3 replicates per time point.

**Supplemental Figure S4.**
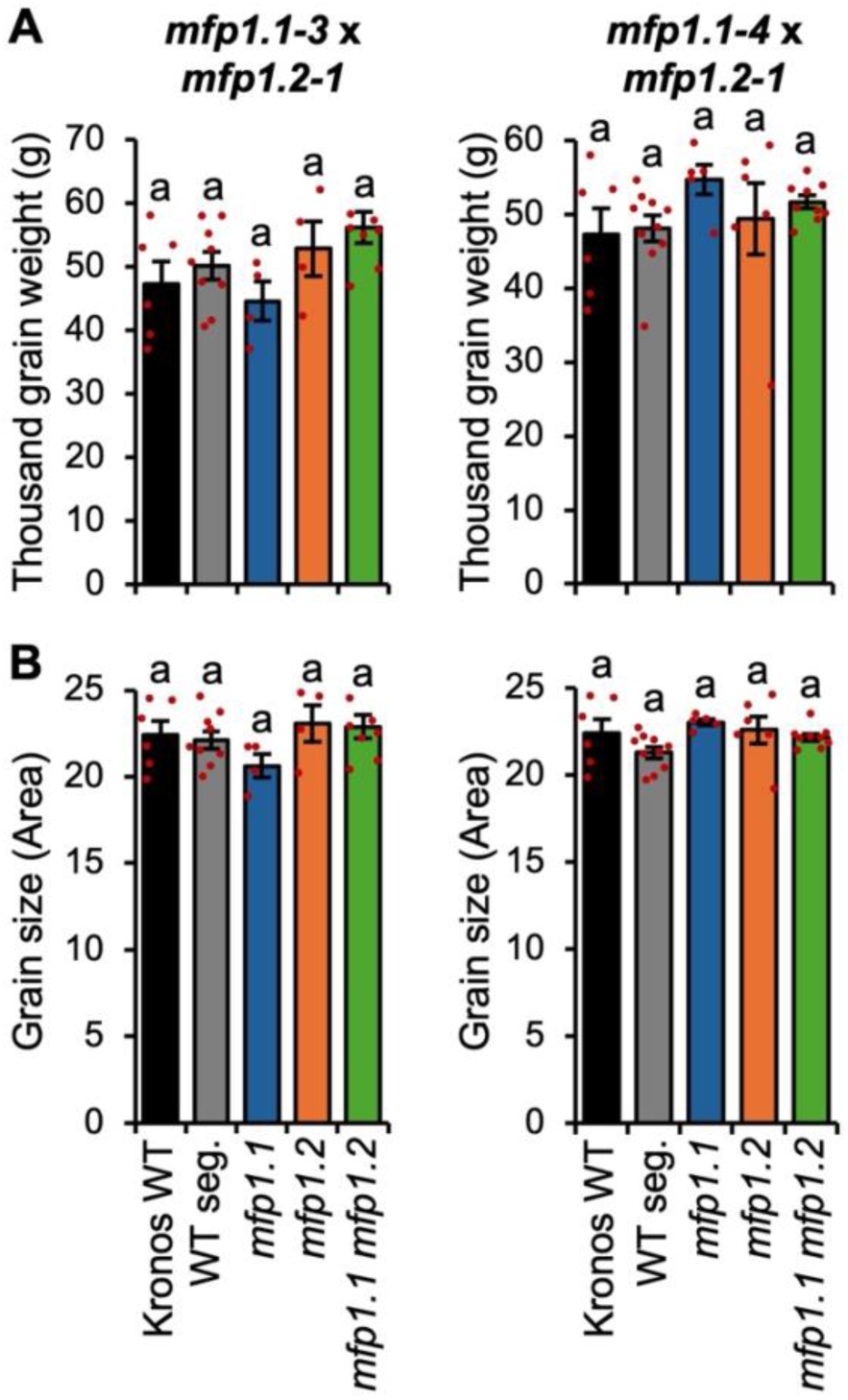
*mfp1.1 mfp1.2* mutants have normal grain phenotypes. The Kronos wild type (WT) was compared with genotypes segregated from both *mfp1.1-3* x *mfp1.2-1* and *mfp1.1-4* x *mfp1.2-1* crosses: including the wild-type segregant control (WT seg.), mutants in both homoeologs of either MFP1.1 (*mfp1.1*) or MFP1.2 (*mfp1.2*), and all homoeologs of both genes (*mfp1.1 mfp1.2*). **A**) Thousand grain weight. **B**) Grain size measured as 2D area. Bars represent the mean ± standard error of the mean from n = 4-10 individual plants, while individual data points are shown in red. Values with different letters are significantly different under a one-way ANOVA and Tukey’s post hoc test at p < 0.05.

**Supplemental Figure S5.**
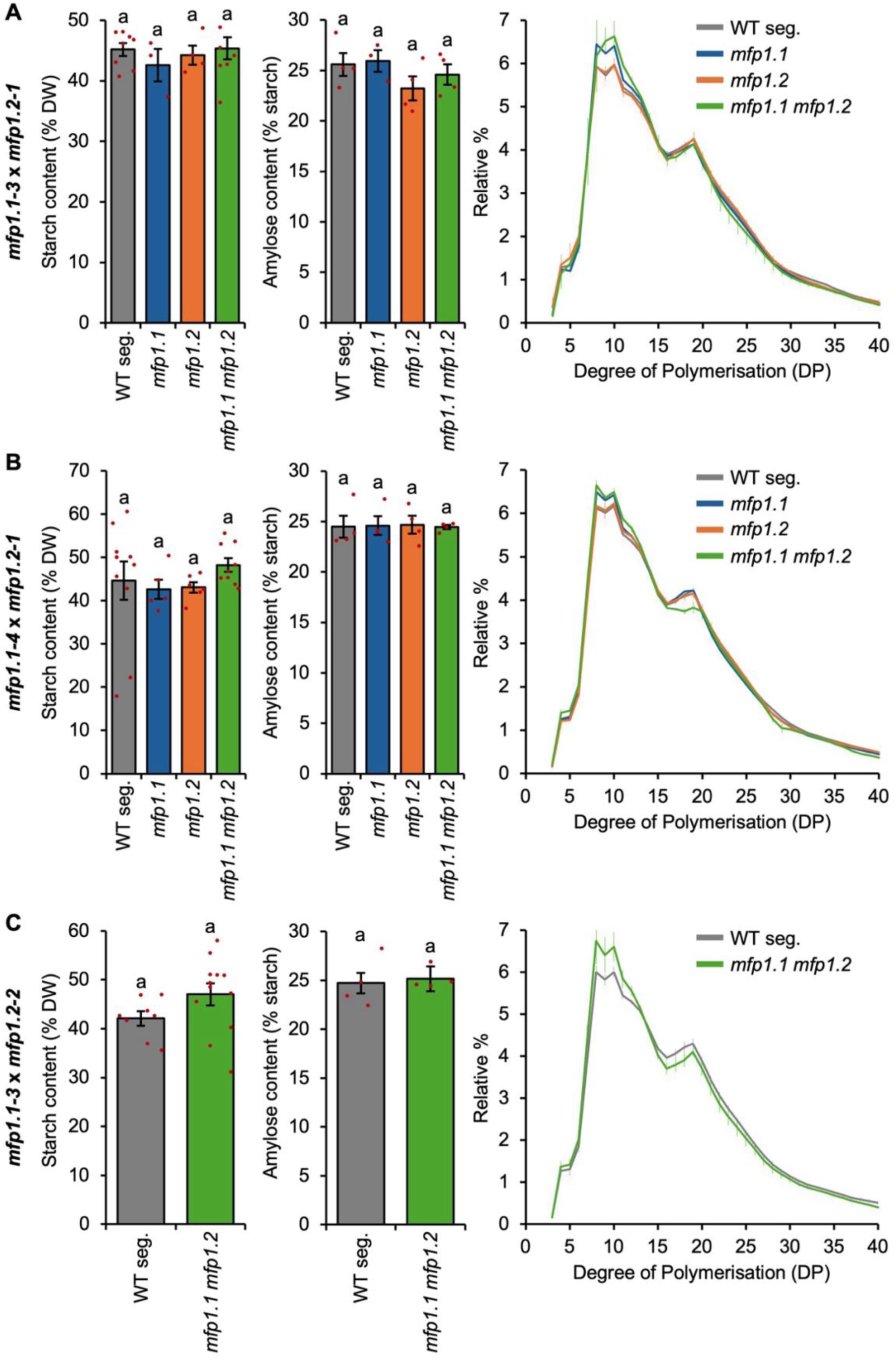
Total starch, amylose, and amylopectin chain length distribution in *mfp1.1 mfp1.2* mutants. The Kronos wild type (WT) was compared with genotypes segregated from the crosses (**A**) *mfp1.1-3* x *mfp1.2-1,* (**B**) *mfp1.1-4* x *mfp1.2-1* and (**C**) for *mfp1.1-3* x *mfp1.2-2*: including the wild-type segregant control (WT seg.), mutants in both homoeologs of either MFP1.1 (*mfp1.1*) or MFP1.2 (*mfp1.2*), and all homoeologs of both genes (*mfp1.1 mfp1.2*). Left panels show total starch content of mature grains, quantified was a percentage dry weight of whole flour. Bars represent the mean ± standard error of the mean (SE) from n = 3-12 individual plants, while individual data points are shown in red. Middle panels show amylose content of purified starch measured using iodine colourimetry. Bars represent the mean ± SE from n = 3-4 individual plants, while individual data points are shown in red. Right panels show amylopectin chain length distribution as determined using High Performance Anion Exchange Chromatography with Pulsed Amperometric Dectection. Values are the mean ± SE on starch extracted from n = 3-4 individual plants. For A and B, values with different letters are significantly different under a one-way ANOVA and Tukey’s posthoc test at p < 0.05. In C, there were no significant differences using a pairwise two-tailed t-test at p < 0.05.

**Supplemental Figure S6.**
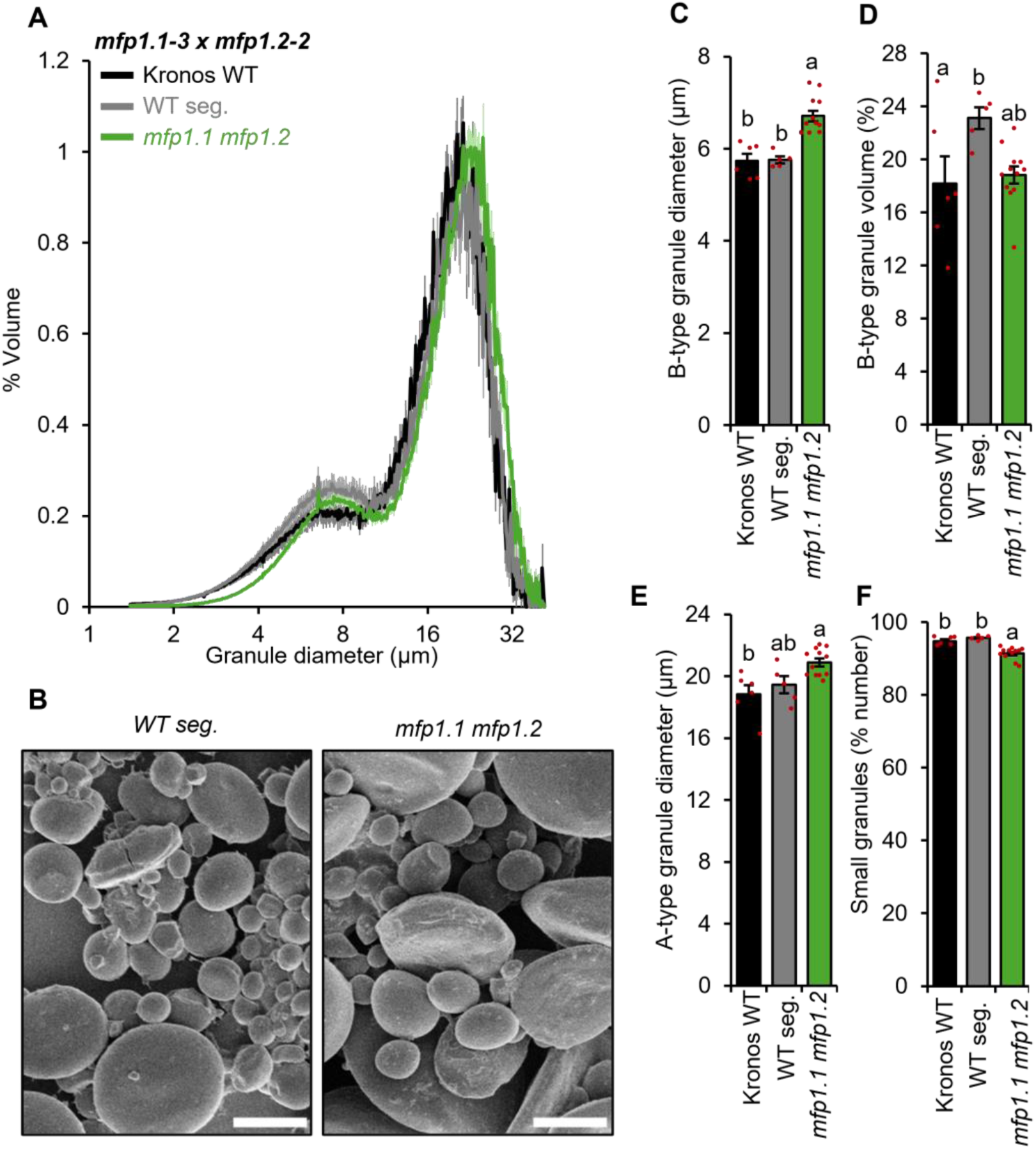
Endosperm starch from the *mfp1.1-3 x mfp1.2-2* mutants have fewer but larger B-type granules. The Kronos wild type (WT) was compared with genotypes segregated from the *mfp1.1-3* x *mfp1.2-2* cross: including the wild-type segregant control (WT seg.), and the mutant in all homoeologs of both MFP1 paralogs (*mfp1.1 mfp1.2*). **A**) Granule size distributions were determined using a Coulter counter, and the data were expressed as relative % volume (of total starch) vs. granule diameter plots. **B**) Scanning electron micrographs of purified endosperm starch. Bars = 10 µm. **C**) Average diameter of B-type granules. **D**) B-type granule volume (% of total starch volume). **E**) Average diameter of A-type granules. In panels **C**-**E**, data were extracted from the relative volume vs. diameter plots in panel **A** by fitting a bimodal mixed normal distribution. **F**) The percentage of small granules by number (defined as number of granules <10 µm, as a percentage of total granule number) was calculated from the Coulter counter data. All plots show the mean from the analysis of n=6-12 replicate starch extractions, each using grains from a separate plant. show the mean from the analysis of n=6-12 replicate starch extractions, each from grains from a separate plant. The shading (panel A) and error bars (panels C-F) represent the standard error of the mean. Values with different letters are significantly different under a one-way ANOVA with Tukey’s post-hoc test (p < 0.05).

**Supplemental Table S1:**
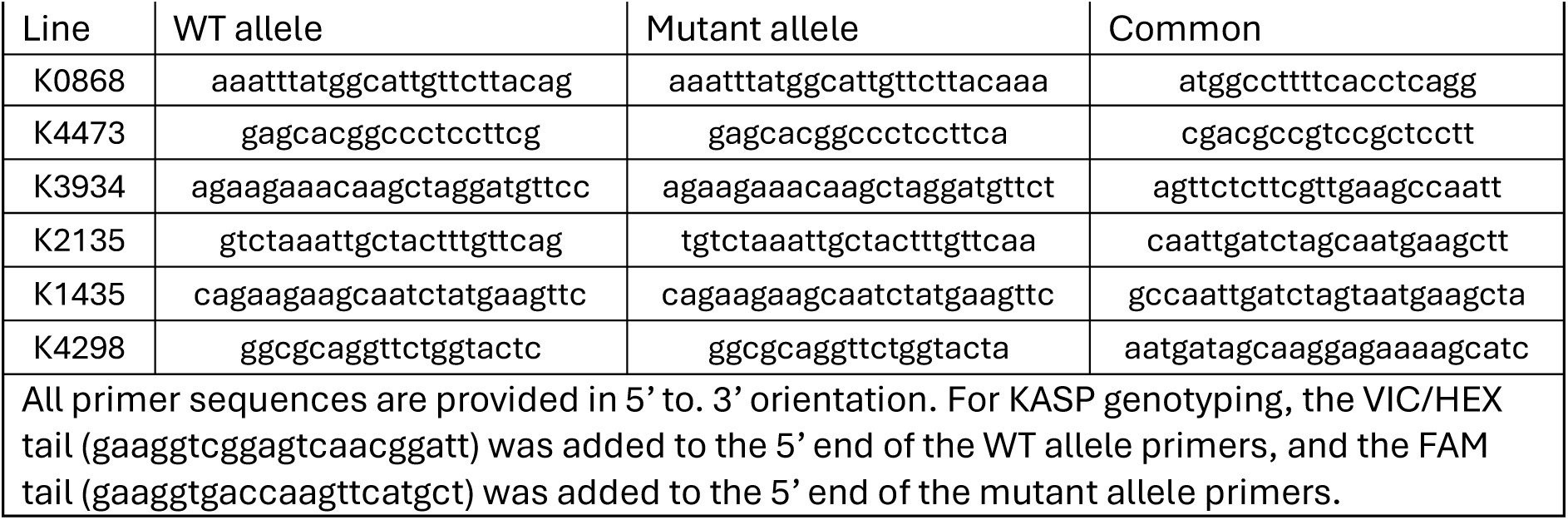
Primers used for KASP genotyping.

